# PsbS confers limited adaptive benefit to C_4_ photosynthesis under fluctuating light

**DOI:** 10.64898/2026.07.09.737394

**Authors:** Russell Woodford, Emma Faraone, Jacinta Watkins, Samuel J. Nix, Susanne von Caemmerer, Robert T. Furbank, Maria Ermakova

## Abstract

Adaptation of plant photosynthesis to dynamic light conditions experienced in natural environments is achieved through specific protective mechanisms. Energy-dependent non-photochemical quenching (qE), regulated by Photosystem II Subunit S (PsbS), is a key process facilitating acclimation to fluctuating light in C_3_ plants, which operate conventional photosynthesis. C_4_ plants, which include some of the world’s most productive and agriculturally important crops, have evolved a distinct high-efficiency photosynthetic pathway. Little is known about the role of specific processes, like qE, in acclimation of C_4_ plants to dynamic light environments. We generated gene-edited lines of the model C_4_ grass *Setaria viridis* lacking PsbS, which were found to be deficient in qE. This deficiency resulted in a modest increase in PSII photoinhibition and a CO_2_ assimilation penalty under light stress in short-term experiments, but photosynthesis and growth under fluctuating light were unaffected. Instead, keeping Photosystem I oxidised through photosynthetic control, negative feedback regulation of the Cytochrome *b*_6_*f* complex, was critical. Therefore, unlike in C_3_ plants, qE does not provide a significant adaptive advantage to C_4_ plants under dynamic light conditions. These findings provide important insights into the biology of C_4_ plants and help prioritise future strategies for improving the productivity and resilience of C_4_ crops.

## Main text

In nature, rapid fluctuations in irradiance caused by sunflecks, canopy movement and cloud cover require continuous re-balancing between the light and dark reactions of photosynthesis^1-4^. This dynamic coordination is necessary to prevent overreduction of the electron transport chain and minimise damage to surrounding photosynthetic proteins, which would otherwise reduce photosynthetic capacity in a phenomenon known as photoinhibition^5,6^. Rapidly-responding photoprotection mechanisms allowing metabolic re-balancing are critical for plant growth under fluctuating light^7-10^. qE, the energy-dependent form of non-photochemical quenching (NPQ), is induced by lumen acidification via the protonation of Photosystem II subunit S (PsbS) and activation of violaxanthin de-epoxidase (VDE), converting the ‘fast’ pool of violaxanthin to zeaxanthin^11-13^. PsbS and zeaxanthin induce conformational changes in the light-harvesting complexes, activating thermal dissipation of excess excitation energy in Photosystem II (PSII) antennae and lowering PSII quantum efficiency^11,12,14^. qE has been extensively studied in C_3_ species, where the lack of PsbS and resulting deficiency in NPQ result in impaired photosynthesis due to increased photoinhibition of PSII^13,15-17^. Consequently, C_3_ species lacking PsbS show impaired growth and reduced seed production under fluctuating light due to photoinhibition caused by an inability to rapidly induce NPQ in response to changes in irradiance^18-22^. Advances in research in C_3_ plants have further identified the importance of qE, recently deciphering the kinetic contributions of various qE components^23,24^ and facilitating the engineering of more responsive qE, increasing productivity of C_3_ crops in naturally fluctuating light^25,26^.

In contrast to C_3_ species, C_4_ plants have evolved a high-efficiency pathway of photosynthesis, with the metabolic C_4_ cycle functioning across mesophyll and bundle sheath (BS) cells^27^. The cycle elevates CO_2_ partial pressure in BS cells where ribulose-1,5-bisphosphate carboxylase/oxygenase (Rubisco) resides, enhancing the carboxylation efficiency of Rubisco while suppressing the oxygenation reaction. This mechanism limits photorespiration in C_4_ plants, which is responsible for up to 50% of crop yield productivity losses in C_3_ plants^28-30^. Overall, this results in an increased productivity of C_4_ plants, including major crops such as maize, sorghum, and sugarcane. The improved productivity and two-cell system of C_4_ photosynthesis, however, present additional considerations for metabolic re-balancing and photoprotection. C_4_ plants have increased energy requirements compared to C_3_, making CO_2_ assimilation more often limited by the energy produced through the light reactions^1,31,32^. This drove C_4_ species to inhabit high light environments and upregulate cyclic electron flow (CEF) around Photosystem I (PSI) via the chloroplast NADH dehydrogenase-like (NDH) complex to increase light energy conversion efficiency^33-35^. Additionally, C_4_ plants are more negatively impacted by fluctuating light than C_3_ species, as, in addition to re-balancing within each cell type, activity of metabolic reactions between mesophyll and BS cells must also be re-balanced each time irradiance changes^36-39^.

Distinct metabolic energy demands have prompted the diversification of chloroplast electron transport chains in mesophyll and BS cells^40^. The two cell types have relatively similar abundance of PSI, while PSII is more abundant in mesophyll cells and NDH is more abundant in BS cells^30,33,41-43^. This enables production of ATP and NADPH through linear electron flow in mesophyll cells and production of predominantly ATP through CEF in BS cells^40^. In line with PSII distribution, PsbS and xanthophyll cycle enzymes are less abundant in BS cells^33,43,44^, suggesting a conserved importance of qE in C_3_ and C_4_ photosynthesis for PSII protection. Indeed, PsbS is necessary for acclimation of maize under extended high light treatments^45^, and natural variation studies have identified genetic control of qE in maize and sorghum under field conditions^46-48^. Despite this, the importance of qE for the adaptation of C_4_ plants to dynamic light environments is still poorly understood.

To identify the importance of qE in adaptation of C_4_ plants to fluctuating light we created two null *PsbS* alleles, *PsbS-1* and *PsbS-2*, in *Setaria viridis* using CRISPR/Cas9 (Fig. S1). Plants with homozygous null alleles (referred to as *psbS-1* and *psbS-2* mutants) lacked PsbS and were deficient in NPQ in rapid light curve analysis, where NPQ should mostly consist of qE^14,49^ (Fig. 1a,b). Instead, mutants had a higher yield of nonregulated non-photochemical reactions in PSII (φ_NO_), but a similar effective quantum yield of PSII (φ_II_), compared to wild-type (WT) plants (Fig. S2). Plants lacking PsbS grown in control conditions with constant daylight of 380 μmol m^−2^ s^−1^ did not differ from WT plants in appearance, biomass, relative chlorophyll content or the maximum quantum efficiency of PSII (F_V_/F_M_) (Fig. 1c-f). Active PSI reaction centers (P_M_) and the yields of photochemical and non-photochemical reactions in PSI were similar between WT and *psbS* plants (Fig. 1g, Fig. S2). No differences in leaf abundance of major thylakoid protein complexes (PSI, PSII, Cytochrome *b*_6_*f* (Cyt*b*_6_*f*), ATP synthase, NDH) or PROTON GRADIENT REGULATION 5 (PGR5) were found between *psbS* and WT plants (Fig. S3). Gas-exchange analysis showed that plants lacking PsbS had lower net CO_2_ assimilation rates at irradiances above 25 *μ*mol m^−2^ s^−1^, compared to WT, which was not due to insufficient intercellular CO_2_ (Fig. 1h,i), but rather a lower maximum electron transport rate (Table S1).

**Fig. 1.**
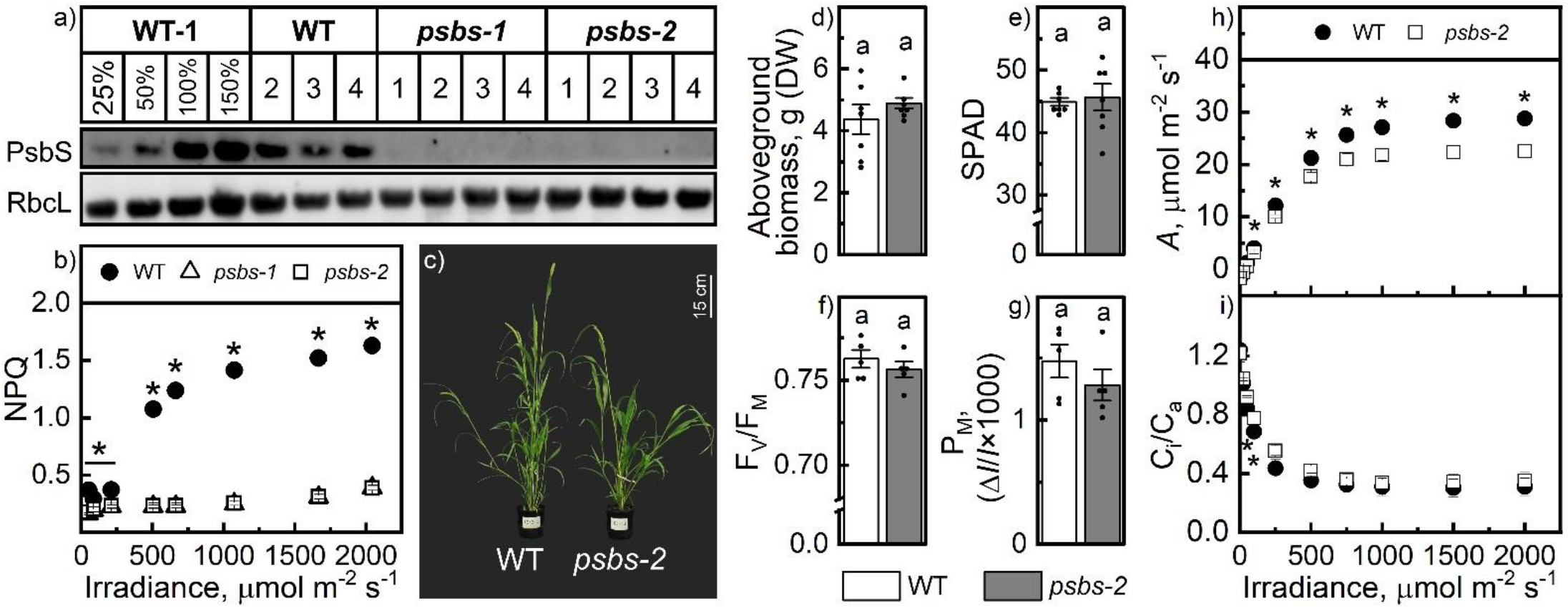
Growth and photosynthesis of *S. viridis* wild type (WT) and plants lacking PsbS (*psbS-1* and *psbS-2*) grown in control conditions, at constant daylight of 380 μmol m^−2^ s^−1^. (**a**) Immunoblotting of total protein extract with antibodies against PsbS and the large subunit of Rubisco (RbcL). 1-4 indicate biological replicates for each genotype. (**b**) Light response of non-photochemical quenching (NPQ). (**c**) Phenotype of plants 5 weeks after germination. (**d**) Aboveground biomass of plants at harvest; DW, dry weight. (**e**) Relative leaf chlorophyll content (SPAD). (**f**) Maximum quantum yield of PSII (F_V_/F_M_). (**g**) Maximum photo-oxidisable P700 (reaction center of PSI), P_M_. (**h, i**) Light response of the net CO_2_ assimilation rate (*A*) and the ratio between intercellular and ambient CO_2_ partial pressures (C_i_/C_a_). Mean ± SE, *n* = 5 biological replicates for (b, f, g), *n* = 7 biological replicates for (d, e), *n* = 4 to 5 biological replicates for (h, i). Asterisks and letters indicate statistically significant differences between genotypes (1-Way ANOVA with Tukey’s post hoc test at *P* < 0.05).

Leaf absorbance at 535 nm, which largely reflects the magnitude of qE^50^, confirmed qE deficiency in *psbS* compared to WT (Fig. 2a). Importantly, this deficiency was not due to changes in thylakoid membrane energization, as evidenced by similar thylakoid proton motive force (*pmf*) in WT and *psbS* (Fig. 2b, Fig. S2). A slower, zeaxanthin-dependent NPQ (qZ) was also unaffected in *psbS* (Fig. 2c), based on xanthophyll content of leaves adapted to dark, control or high irradiance for 30 min^14,49^. Interestingly, the lack of qE had little effect on instantaneous responses of net CO_2_ assimilation rates to fluctuating light (repeated cycles of 3 min at 2000 *μ*mol m^−2^ s^−1^ and 3 min at 500 *μ*mol m^−2^ s^−1^) (Fig. 3a). Again, *psbS* showed no differences in PSI photochemical efficiency (Fig. S4), but had similar to WT φ_II_, consistently lower NPQ and consistently higher φ_NO_; however, NPQ in the mutant slowly increased over the course of the experiment (Fig. 3b-d). Notably, after the fluctuating light sequence, relaxation of φ_NPQ_ was observed in the dark in WT but not in *psbS* (Fig. 4c), which was consistent with almost twice the loss in PSII maximum efficiency (Fig. 3b inset), indicating increased photoinhibition of PSII in the absence of qE^14,51^.

**Fig. 2.**
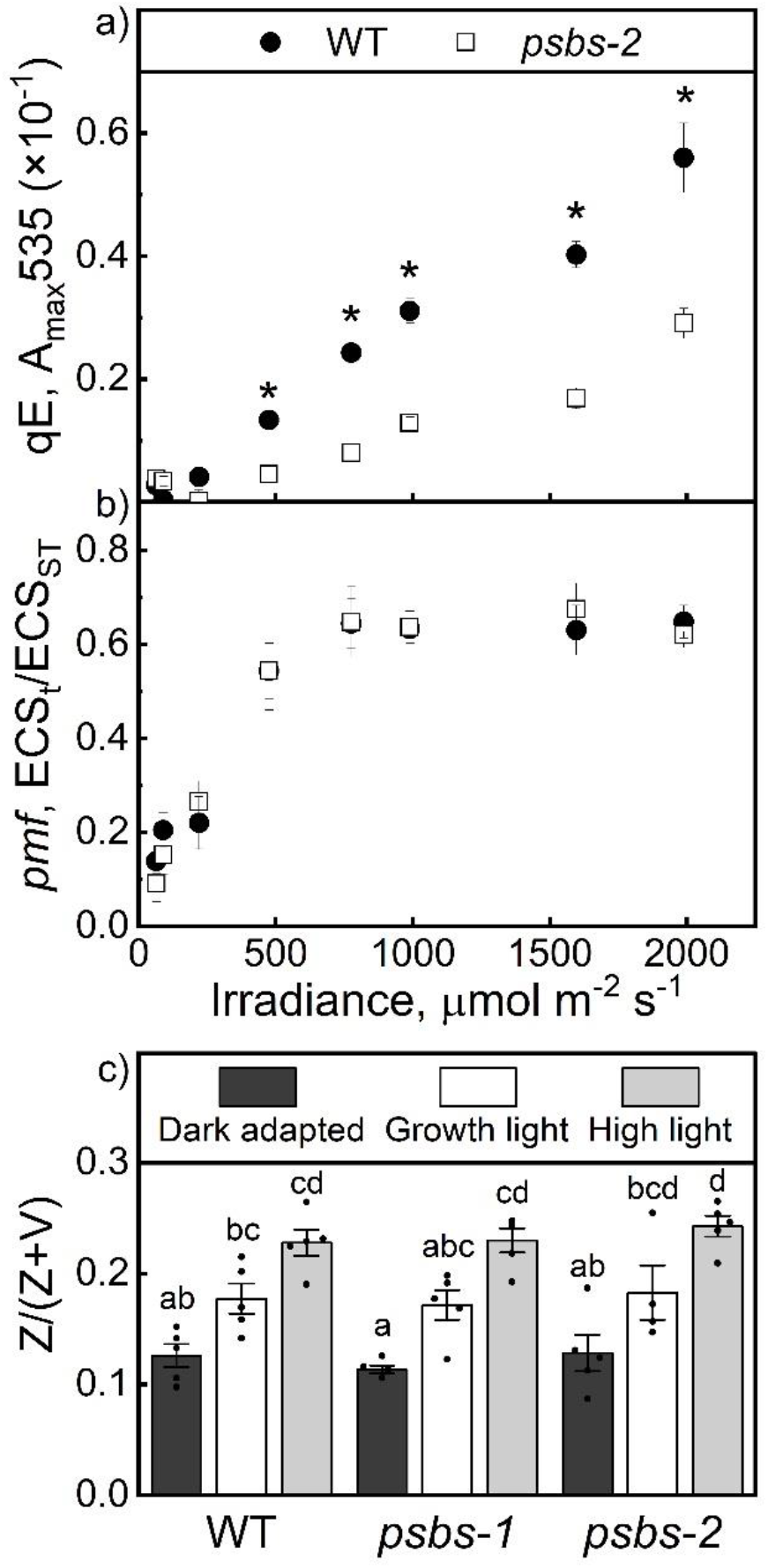
Non-photochemical quenching and thylakoid membrane energisation analyses in wild-type (WT) *S. viridis* and plants lacking PsbS (*psbS-1* and *psbS-2*) grown in control conditions. (**a**) Maximum leaf absorbance at 535 nm, indicative of energy-dependent NPQ (qE), at different irradiances. (**b**) Proton motive force (*pmf*) at different irradiances. (**c**) Relative abundance of xanthophyll cycle pigments in leaves adapted for 30 min to different irradiances (growth light, 380 μmol m^-2^ s^-1^; high light 2000 μmol m^-2^ s^-1^); Z, zeaxanthin; V, violaxanthin. Mean ± SE, *n* = 4 to 5 biological replicates in (a,c). *n* = 5 biological replicates in (b). Asterisks indicate statistically significant differences between genotypes for (a, b) (1-Way ANOVA with Tukey’s post hoc test at *P* < 0.05). Letters indicate significant differences between genotypes and conditions for (c) (2-Way ANOVA with Tukey’s post hoc test at *P* < 0.05).

**Fig. 3.**
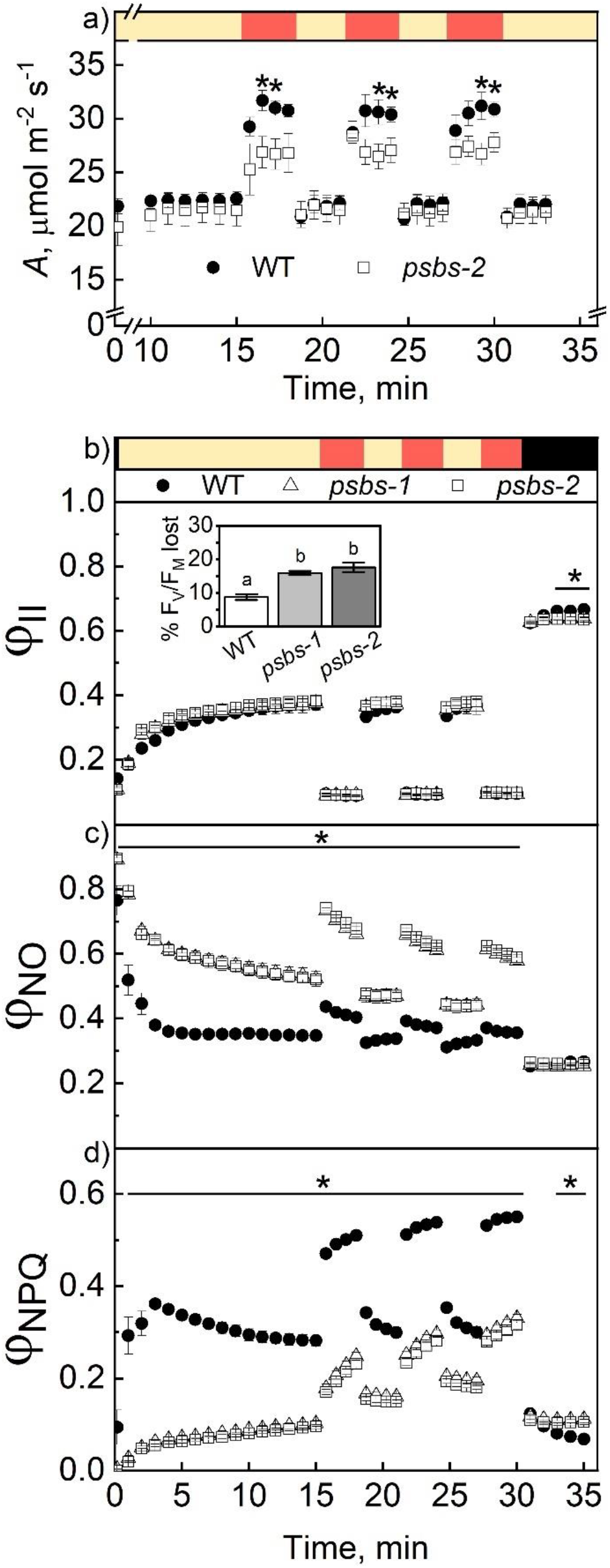
Effects of fluctuating light on wild-type (WT) *S. viridis* and plants lacking PsbS (*psbS-1* and *psbS-2*). Plants were grown in control conditions and subjected to fluctuating light: yellow bars, 500 μmol m^−2^ s^−1^; red bars, 2000 μmol m^−2^ s^−1^; black bars, darkness. (**a**) Net CO_2_ assimilation rate (*A*). (**b**) The effective quantum yield of PSII (φ_II_). (**b, inset**) Percentage loss of maximum quantum yield of PSII (F_V_/F_M_) following the fluctuating light experiment. (**c**) The yield of nonregulated non-photochemical reactions (φ_NO_). (**d**) The yield of NPQ (φ_NPQ_). Mean ± SE, *n* = 5 biological replicates. Bars with asterisks and letters indicate significant differences between both mutants and WT (1-Way ANOVA with Tukey’s post hoc test at *P* < 0.05).

**Fig. 4.**
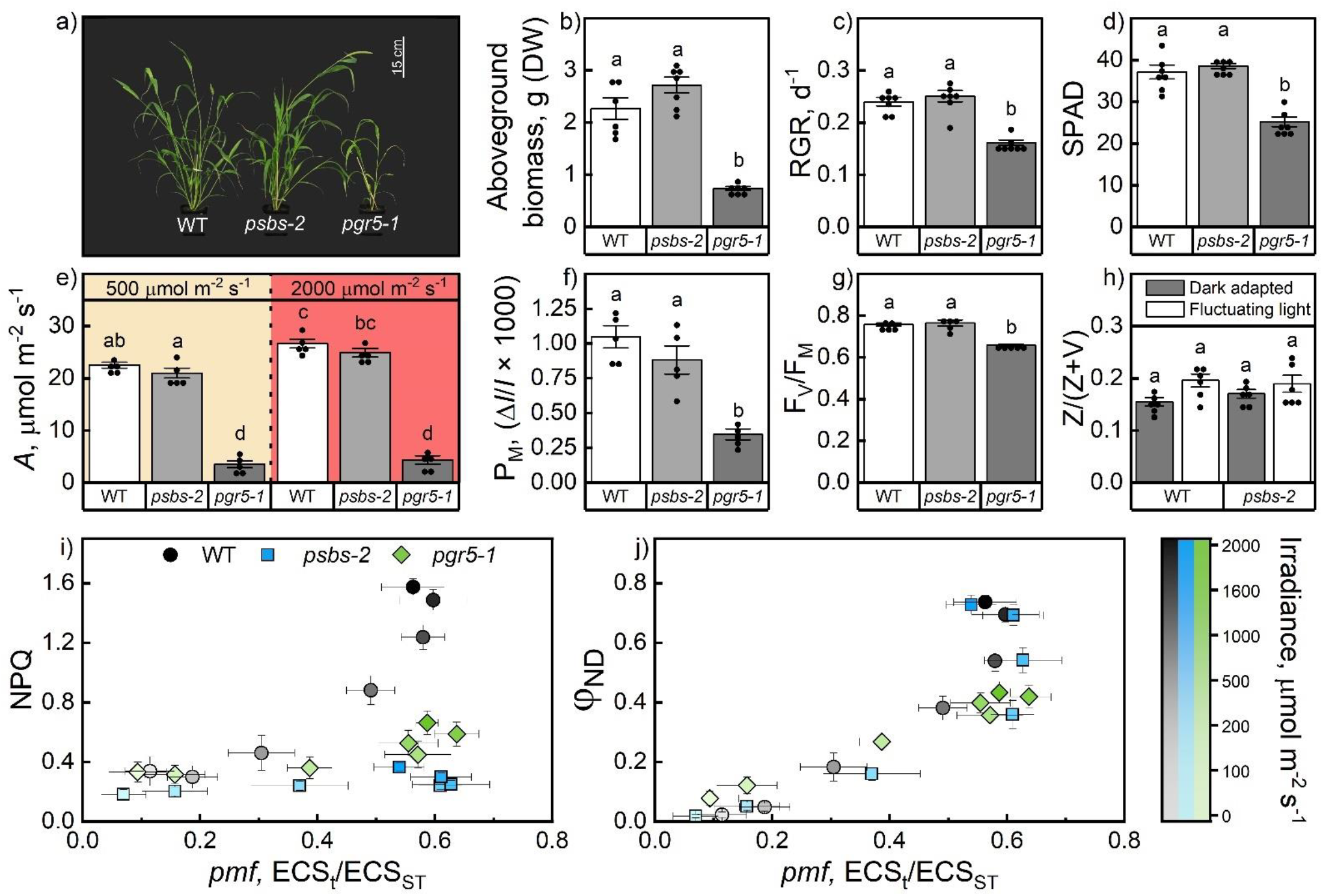
Growth and photosynthesis of *S. viridis* wild type (WT) and plants lacking PsbS (*psbS-2*) or PGR5 (*pgr5-1*) grown at fluctuating daylight (repeating cycles of 5 min at 250 μmol m^−2^ s^−1^ and 1 minute at 1000 μmol m^−2^ s^−1^). (**a**) Phenotype of plants 5 weeks after germination. (**b**) Aboveground biomass at harvest; DW, dry weight. (**c**) Relative growth rate (RGR) based on 3D leaf area. (**d**) Relative leaf chlorophyll content (SPAD). (**e**) Net CO_2_ assimilation rate (*A*) at 500 and 2000 μmol m^−2^ s^−1^ irradiance. (**f**) Maximum photo-oxidisable P700 (reaction center of PSI), P_M_. (**g**) Maximum quantum yield of PSII (F_V_/F_M_). (**h**) Relative abundance of xanthophyll cycle pigments in dark and light-adapted leaves; Z, zeaxanthin; V, violaxanthin. (**i**) The relationship between non-photochemical quenching (NPQ) and proton motive force (*pmf)* at corresponding irradiances. (**j**) The relationship between the non-photochemical yield of PSI due to donor side limitation (φ_ND_) and *pmf* at corresponding irradiances. Mean ± SE, *n* = 6 to 7 biological replicates in (b), *n* = 7 biological replicates in (c,d), *n* = 5 biological replicates in (e-g), *n* = 6 biological replicates in (h). Letters indicate significant differences between genotypes for (b, c, d, f, g) (one-way ANOVA with Tukey’s post hoc test at *P* < 0.05). Letters indicate significant differences between genotypes and conditions for (e, h) (two-way ANOVA with Tukey’s post hoc test at *P* < 0.05).

An absence of PGR5, diminishing both qE and photosynthetic control, negative feedback on plastoquinol oxidation at Cyt*b*_6_*f* that restricts electron flow to PSI, was recently shown to be detrimental for the acclimation of *S. viridis* to fluctuating light^52^. To identify whether a lack of qE alone would hinder acclimation to fluctuating light, we grew plants under similar conditions: repeated cycles of 5 min at 250 *μ*mol m^−2^ s^−1^ and 1 min at 1000 *μ*mol m^−2^ s^−1^ throughout the day. The biomass of WT plants was only about 50% when grown at fluctuating light compared to constant daylight, but the biomass and relative growth rate of *psbS* and WT grown under fluctuating light were identical (Fig. 1c-d, Fig. 4a-c). Furthermore, no differences in relative chlorophyll content, CO_2_ assimilation rates, P_M_, F_V_/F_M_ or zeaxanthin content were found between *psbS* and WT plants grown at fluctuating light (Fig. 4d-h). This was in sharp contrast with the *pgr5* mutant (Fig. 6a-g), which was severely affected by fluctuating light but grew similar to WT under constant daylight^52^. Comparison of photochemical and non-photochemical yields of PSII and PSI (Fig. S5) with thylakoid membrane energisation parameters (Fig. S6) identified that the difference in fluctuating light adaptation between *psbS* and *pgr5* could be attributed to a lack of photosynthetic control in *pgr5*, seen as an inability to oxidise PSI at high irradiances (low φ_ND_), even at WT-levels of *pmf* (Fig. 4i-j).

Our results show that the function of qE in safe dissipation of excess excitation energy to decrease PSII photoinhibition is similar between C_3_ and C_4_ plants, consistent with previous literature^15-17,45,53^. The lack of qE did result in increased susceptibility to PSII photoinhibition in plants grown at constant light, leading to decreased CO_2_ assimilation under high light and during the high light phases of fluctuating light (Fig. 1h, Fig. 3a). The *psbS* mutant dissipates excess energy at PSII through φ_NO_, constitutive thermal dissipation via reaction centre quenching, which provides some photoprotection but is more likely to cause PSII photoinhibition than qE^54^. Avoiding φ_NO_ and PSII photoinhibition is one of the priorities of C_3_ plants, as seen through the reduced growth and increased PSII photoinhibition of C_3_ *psbS* mutants grown under fluctuating light^18-22^. Interestingly, growth and photosynthesis of *S. viridis psbS* was not affected under fluctuating light (Fig. 4), indicating a markedly different approach to managing photoinhibition and photoprotection in C_4_ plants.

While *pgr5* lacks both qE and photosynthetic control, direct comparison with *psbS* indicated the lack of photosynthetic control is primarily responsible for the severe fluctuating light phenotype (Fig. 4). Photosynthetic control and qE have photosystem-specific photoprotective outputs: while qE helps to avoid photoinhibition of PSII, photosynthetic control keeps PSI oxidised, allowing safe charge recombination to prevent PSI photoinhibition^53,55,56^. Compared with PSII photoinhibition, which can be remedied relatively rapidly due to the dedicated repair cycle at the cost of ATP, PSI photoinhibition is more detrimental because recovery requires reassembly of the entire complex over timescales of days to weeks^6,57-60^. C_4_ photosynthesis relies heavily on CEF around PSI for ATP production^33,61^. Provided PSI remains intact and runs CEF, absence of PsbS can be overcome via PSII repair without significantly impairing electron transport^62,63^. However, NDH-CEF per se did not confer a specific benefit to C_4_ photosynthesis under fluctuating light (Fig S7), which is in contrast to C_3_ plants^64^ and further highlights the primary importance of NDH-CEF for ATP generation independently of conditions. Instead, keeping PSI intact through PGR5 emerges as the most critical for adaptation to fluctuating light. Therefore, by increasing the abundance of PGR5, but not PsbS, compared to C_3_ plants^34,35,65^, and distributing PGR5 equally between the cell types^33,43,66^, C_4_ plants prioritise photosynthetic control and photoprotection of PSI in both cell types.

Increasing evidence suggests that photosynthetic control is not simply an intrinsic property of Cyt*b*_6_*f* strictly responding to lumen pH but is subject to complex regulation (reviewed in Degen and Johnson ^67^). Our results support this concept in C_4_ plants: in the absence of PGR5, photosynthetic control could not be established even by the WT-levels of *pmf* under fluctuating light (Fig. 4j). As the relative importance of photosynthetic control over qE for acclimation to fluctuating light appears even greater in C_4_ than C_3_ plants, increasing photosynthetic efficiency and productivity of C_4_ crops in dynamic light environments will require a deeper understanding of the molecular mechanisms governing this process. These findings therefore highlight an urgent need to elucidate the regulation of photosynthetic control, many aspects of which remain poorly understood.

## Materials and Methods

### CRISPR-Cas9 mutant generation

*S. viridis* cv. ME034V-1 plants with null *psbS-1* and *psbs-2* alleles were generated using CRISPR/Cas9 gene-editing as described in Ermakova, et al. ^33^, using the *PsbS* sequence obtained from Phytozome (Sevir.5G400800). Cas9 was targeted to the second and third exons of *PsbS* using two gRNAs, selected with CRISPOR^68^, which were assembled into a synthetic polycistronic gene according to Xie, et al. ^69^. The construct was assembled using the Golden Gate cloning system^70^ and transformed into *S. viridis* tissue culture using *Agrobacterium tumefaciens* strain AGL1 following the procedure described in Osborn, et al. ^71^. Two null alleles of *PsbS* were identified by sequencing, and the progenies of T_1_ plants lacking T-DNA and homozygous for null alleles were used in all experiments (Fig. S1).

### Plant growth conditions

*S. viridis* plants were grown in 0.5 L pots with seed and cutting mix (Debco, Tyabb, Australia or Scotts, Bella Vista, Australia) and 5 g L^-1^ of Osmocote slow-release fertilizer (Scotts, Bella Vista, Australia). Plants were grown in controlled environment chambers (Conviron, Winnipeg, Canada) with ambient CO_2_, 28°C (day) and 22°C (night), and 60% humidity, and either a constant (380 μmol m^−2^ s^−1^, 16 h photoperiod) or a fluctuating daylight. The fluctuating daylight cycle consisted of 5 min at 250 μmol m^−2^ s^−1^ and 1 minute at 1000 μmol m^−2^ s^−1^ repeating for 16 h and providing a total integrated irradiance of 375 μmol m^−2^ s^−1^. All plants were initially germinated under constant daylight. Two weeks post-germination, half of the plants were transferred to the fluctuating daylight treatment in a separate growth cabinet, and half were retained under constant daylight. Concurrent growth experiments make the results in Fig. 1 and Fig. 4 directly comparable. All measurements and analyses were performed on the youngest fully expanded leaves of three-to-four-week-old plants.

### Plant phenotyping

3D plant imaging was performed using a PlantEye F600 Multispectral 3D scanner (Phenospex, Heerlen, The Netherlands). Whole-plant imaging generated 3D point clouds, with 3D leaf area calculated using voxel-based reconstructions. Relative growth rate was calculated from 3D leaf area (LA) as: 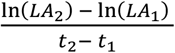, using measurements performed 14 and 28 days after sowing. The total aboveground dried biomass was determined on 45-day-old *S. viridis* plants after oven-drying plant material at 80°C for one week. Photosynthesis at ambient irradiance and relative leaf chlorophyll content (SPAD) were analysed with MultispeQ V2.0 (PhotosynQ, East Lansing, MI, USA) using the RIDES 2.0 protocol^72^. For plants growing under fluctuating light, measurements were performed during the low irradiance periods.

### Leaf spectroscopic and fluorescence analyses

To measure light responses of photosystems, simultaneous measurements of chlorophyll fluorescence and P700 redox spectroscopy were performed using a DUAL-KLAS-NIR or DUAL-PAM/F (Heinz Walz, Effeltrich, Germany) under ambient air in well-ventilated conditions, but without controlling gas flow. DUAL-KLAS-NIR measurements were performed under red actinic light (635 nm) with red light saturating pulses (635 nm, 300 ms, 12,000 μmol m^-2^ s^-1^), with P700 redox state determined from near-infrared signal deconvolution using differential model plots for WT *S. viridis*^73-75^. DUAL-PAM/F measurements were performed under red actinic light (635 nm) with red light saturating pulses (635 nm, 300 ms, 12,000 μmol m^-2^ s^-1^), with P700 redox state assessed using a dual wavelength unit (830/875 nm). Leaves were dark-adapted for 30 min before all measurements.

For DUAL-KLAS-NIR measurements, the maximal P700^+^ signal (P_M_) was determined using the pre-installed NIRmax routine^73-75^. Subsequently, a saturating pulse was applied to determine the maximum (F_M_) and minimum (F_0_) levels of fluorescence, with the maximum quantum yield of PSII (F_V_/F_M_) calculated as (F_M_ - F_0_) / F_M_. For Dual-PAM/F measurements, first F_M_, F_0_ and F_V_/F_M_ were determined as describe above. Next, the maximal P700^+^ signal (P_M_) was recorded by applying a saturating pulse at the end of an 8-s far-red light (720 nm) illumination, with the minimal P700^+^ signal (P_0_) recorded directly following the saturating pulse. Leaves were then acclimated to 500 μmol m^−2^ s^−1^ for 15 minutes.

Light response curves were recorded by progressively increasing the actinic light from 0 to 2000 μmol m^−2^ s^−1^ and applying a saturating pulse at the end of every 120-s illumination period. Upon the application of each saturating pulse, F (the steady-state fluorescence level) and F_M_’ (the maximum level of fluorescence under light) were recorded. The steady-state P700^+^ signal (P) and the maximal P700^+^ signal under light (P_M_’) were similarly recorded prior to and upon the application of each saturating pulse, respectively. This enabled the monitoring of the partitioning of absorbed light energy within PSII between the photochemical [φ_II_ = (F_M_’ – F) / F_M_] and non-photochemical reactions, including the regulated [φ_NPQ_ = (F_M_ – F_M_^‘^) / F_M_] and non-regulated (φ_NO_ = F / F_M_) fractions^76^. Similarly, the photochemical yield of PSI [φ_I_ = (P_M_’ – P) / (P_M_ – P_0_)], the non-photochemical yield of PSI due to acceptor side limitation [φ_NA_ = (P_M_ – P_M_’) / (P_M_ – P_0_)] and the non-photochemical yield of PSI due to donor side limitation [φ_ND_ = (P – P_0_) / (P_M_ – P_0_)] were calculated as described in Klughammer and Schreiber ^77^ and Schreiber and Klughammer ^75^.

For fluctuating light experiments leaves were subjected to an alternating light sequence, with actinic irradiance oscillating between 2000 μmol m^-2^ s^-1^ and 500 μmol m^-2^ s^-1^ at 3-minute intervals. Following the third high light period, actinic light was ceased for 5 mins while recording continued. Saturating pulses were applied every 45 seconds throughout the measurement and the photochemical yields of PSI and PSII were calculated as described above.

### Thylakoid membrane energisation

ECS signal was monitored with the Dual PAM-100 (Heinz Walz) equipped with the P515/535 emitter-detector module. ECS was estimated from the absorbance changes between 550 and 520 nm and normalised for the amplitude of ECS response to a saturating pulse (20 μs, 14,000 μmol m^−2^ s^−1^) measured from the dark-adapted leaves (ECS_ST_). As the Dual PAM-100 uses LED illumination, rather than laser, ECS normalisation using a true single turnover pulse (<1 μs) is not possible^78^. While normalising ECS with a >1 μs pulse is inappropriate for determining electron flow rates or transmission coefficients, normalising with a pulse length up to 50 μs is routine for comparative studies^67,79,80^. Changes in ECS signal were measured during 3 min illumination periods of increasing irradiance. After each illumination period, the light was switched off for 20 s and the kinetics of ECS signal decay were analysed to determine proton motive force (*pmf*), proton conductivity of the thylakoid membrane (*g*_H_+), and the light-driven proton flux across the thylakoid membrane (*v*_H_+). *Pmf* was estimated as the total change in amplitude of ECS signal (ECS_t_) normalised for ECS_ST_^81^, *g*_H_+ was calculated as the inverse time constant of the first-order exponential decay during the first 150 ms following light termination^82^, and *v*_H_+ was calculated as the linear rate of ESC decay during the first 30 ms following light termination^83^.

Leaf absorbance signal at 535 nm, informative of qE^50^, was measured simultaneously with ECS response during the experiment described above. At each irradiance, the highest 535 nm absorbance signal (A_max_535) was determined as the difference between the average 535nm absorbance signal during the final 5 seconds of the actinic light period and the average 535 nm absorbance signal during the final 5 seconds of the preceding dark period. The resulting A_max_535 signals for each irradiance were then fit to a linear regression, with A_max_535 values then baseline corrected to the zero intercept.

### Gas-exchange analysis

Gas-exchange analysis was performed using a portable gas-exchange system LI-6800 (LICOR, Lincoln, Nebraska, USA) equipped with a 6800-01A fluorometer head. For all measurements 90% red/10% blue actinic light was used with 25°C leaf temperature, 55% relative chamber humidity, flow rate of 500 μmol s^−1^, flow pressure of 0.1 kPa and fan speed of 10,000 RPM. For light response curves, leaves were first equilibrated at 500 μmol mol^-1^ CO_2_ on the reference side and 500 μmol m^-2^ s^-1^ irradiance for 15 minutes. After equilibration, CO_2_ assimilation was recorded at irradiances increasing from 0 to 2000 μmol m^-2^ s^-1^ at 3-minute intervals. Light response curves were fit using the biochemical model of C_4_ photosynthesis parameterised for *S. viridis*^84,85^. For gas-exchange analysis under a fluctuating light regime, after equilibration, plants were subjected to a fluctuating light sequence (repeating cycles of 3 min at 2000 *μ*mol m^−2^ s^−1^ and 3 min at 500 *μ*mol m^−2^ s^−1^), with data logged every 45 s.

### Xanthophyll analysis

Leaf discs for xanthophyll analysis were sampled in the middle of the day during the low-light phase (250 *μ*mol m^−2^ s^−1^) of the fluctuating daylight sequence, flash-frozen in liquid nitrogen and freeze-dried. Pigments were extracted using acetone-ethyl acetate (6:4, v/v) as described in Alagoz, et al. ^86^, and 100 *μ*g of α-tocopheryl acetate (Sigma) was added to each sample as an internal standard prior to separation. Pigments were separated on a high-performance liquid chromatograph (Agilent 1200, Agilent Technologies) using a C30 column (4.6 × 250 mm; YMC) coupled to a 5 *μ*m C30 guard column (4.0 × 23 mm). Forty microliters sample injections were separated at a flow rate of 1 mL min^−1^ with mobile phases of 100% methanol and 100% methyl tert-butyl ether, with a gradient elution performed as described in Alba, et al. ^87^.

### Immunoblotting

Protein extraction from leaf discs, SDS-PAGE, and immunoblotting was performed as described in Ermakova, et al. ^88^. Samples were normalised on leaf area basis, separated by SDS-PAGE and transferred to nitrocellulose membrane. Samples were probed with antibodies against key photosynthetic proteins: PGR5 (1:3000, AS163985, Agrisera, Vännäs, Sweden), NdhH (NDH, 1:3000, AS164065, Agrisera), Rieske (Cyt*b*_6_*f*, 1:5000, AS08330, Agrisera), RbcL (Rubisco, Martin-Avila, et al. ^89^), PsaB (PSI, 1:5000, AS10695, Agrisera), AtpB (ATP synthase, 1:10,000, Agrisera), and PsbS (1:3000, AS09533, Agrisera), Lhcb2 (1:10,000, AS01003; Agrisera), PEPC (1:10 000, Karki, et al. ^90^). Immunoblots were imaged and quantified using ChemiDoc Go Imaging System and Image Lab software (Bio-Rad, Hercules, CA, USA).

### Statistical analysis

One-way and two-way ANOVAs were performed in OriginPro 2023b using a significance threshold of *P* < 0.05. Details of replication, post-hoc tests, and *P*-values are provided in the figure legends.

## Acknowledgements

We thank Dr Xueqin Wang for help with *S. viridis* transformation. We acknowledge the use of the facilities, and scientific and technical assistance of the Australian Plant Phenomics Network (APPN), which is supported by the Australian Government’s National Collaborative Research Infrastructure Strategy (NCRIS). This work was supported by a Postgraduate Internship Award from APPN to RW.

## Authors contribution

ME and RW designed research; ME, SvC and RTF supervised research; RW, EF, JW and SJN performed experiments; RW, EF, JW and ME analysed data; RW, EF and ME wrote the manuscript; all authors discussed results and edited the final manuscript.

