## Supplementary materials for "PsbS confers limited adaptive benefit to C_4_ photosynthesis under fluctuating light"

**Table S1.** Photosynthetic parameters of wild type (WT) and PsbS-deficient (*psbS-2*) *S. viridis* plants fit from light response ($A/I$) curves of $CO_2$ assimilation (Fig. 1g). $R_d$, day respiration rate; $J_{max}$, maximum linear electron transport rate; $J_{max\ cyc}$, maximum cyclic electron transport rate; $J_{max\ 1}$, maximum electron transport rate through PSI; $\theta$, empirical curvature factor; $\phi_{CO2}$, quantum yield of $CO_2$ assimilation; $\Gamma_{light}$, light compensation point (see details in Woodford, et al. [85]. Asterisks indicate statistically significant differences between the *psbs*-2 and WT (1-Way ANOVA with Tukey's post hoc test at $P < 0.05$). Mean ± SE, $n$ = 4 to 5 biological replicates.

| **Parameter** | **WT** | ***psbs-2*** |
|---|---|---|
| $R_d$ (µmol m$^{-2}$ s$^{-1}$) | 0.96 ± 0.12 | 1.67 ± 0.08* |
| $J_{max}$ (µmol m$^{-2}$ s$^{-1}$) | 139.46 ± 3.67 | 116.98 ± 5.40* |
| $J_{max\ cyc}$ (µmol m$^{-2}$ s$^{-1}$) | 114.11 ± 3.00 | 95.71 ± 4.42* |
| $J_{max\ 1}$ (µmol m$^{-2}$ s$^{-1}$) | 253.57 ± 6.67 | 212.69 ± 9.82* |
| $\theta$ | 0.92 ± 0.02 | 0.89 ± 0.02 |
| $\phi_{CO2}$ | 0.057 ± 0.004 | 0.049 ± 0.003 |
| $\Gamma_{light}$ (µmol m$^{-2}$ s$^{-1}$) | 28.80 ± 0.88 | 36.32 ± 0.79* |

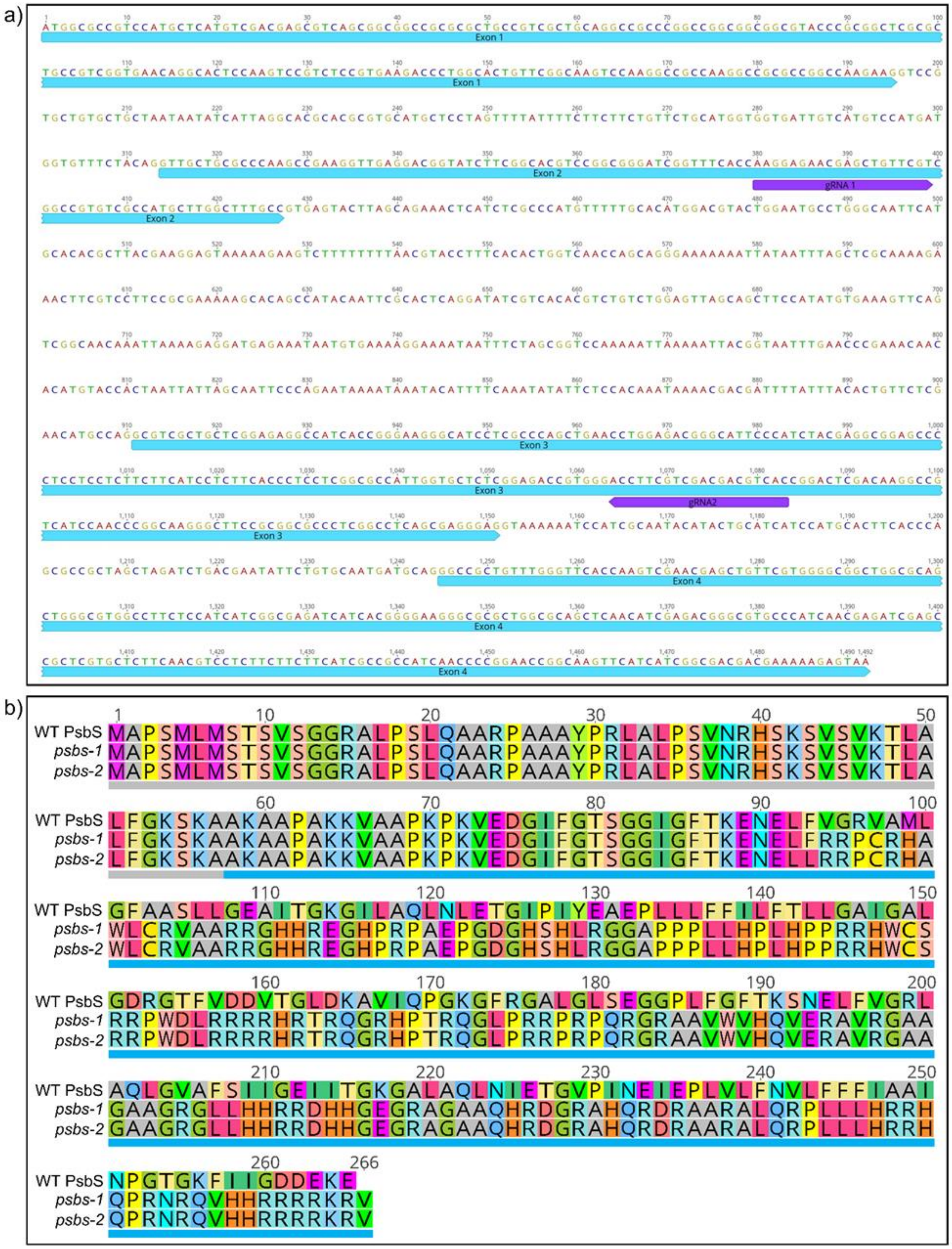


**Fig. S1.** Two new *PsbS* alleles obtained via CRISPR/Cas9 genome-editing of *S. viridis*. (**a**) Location of guide RNAs (gRNAs) targeting the second and third exons of the *PsbS* genomic sequence. (**b**) Amino acid sequences of PsbS and proteins predicted to be encoded by the new alleles. Grey bar underneath indicates chloroplast signal peptide, blue bar – mature protein.

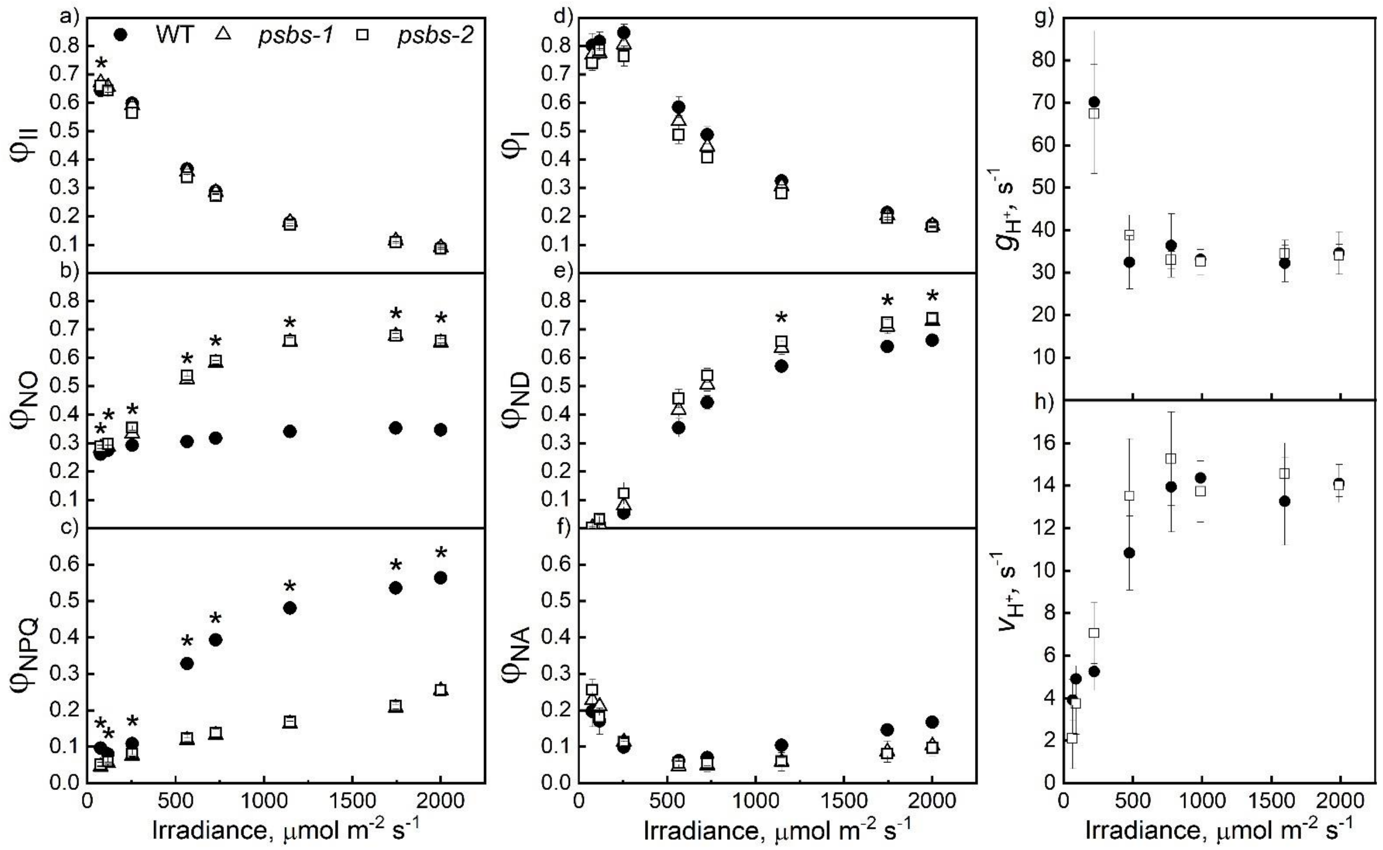


**Fig S2**. Light response of quantum yields within PSII (**a** to **c**) and PSI (**d** to **f**), and thylakoid membrane energisation parameters (**g**, **h**) in wild-type (WT) *S. viridis* and plants lacking PsbS (*psbS-1* and *psbS-2*) grown in control conditions. (**a**) The effective quantum yield of PSII ($\varphi_{II}$). (**b**) The yield of nonregulated non-photochemical reactions ($\varphi_{NO}$). (**c**) The yield of NPQ ($\varphi_{NPQ}$). (**d**) The photochemical yield of PSI ($\varphi_{I}$). (**e**) The non-photochemical yield of PSI due to donor side limitation ($\varphi_{ND}$). (**f**) The non-photochemical yield of PSI due to acceptor side limitation ($\varphi_{NA}$). (**g**) Proton conductivity of the thylakoid membrane ($g_{H}$+). (**h**) The light-driven proton flux across the thylakoid membrane ($v_{H}$+). Mean ± SE, $n$ = 5 biological replicates. Asterisks indicate significant differences between both mutants and WT (1-Way ANOVA with Tukey's post hoc test at $P < 0.05$).

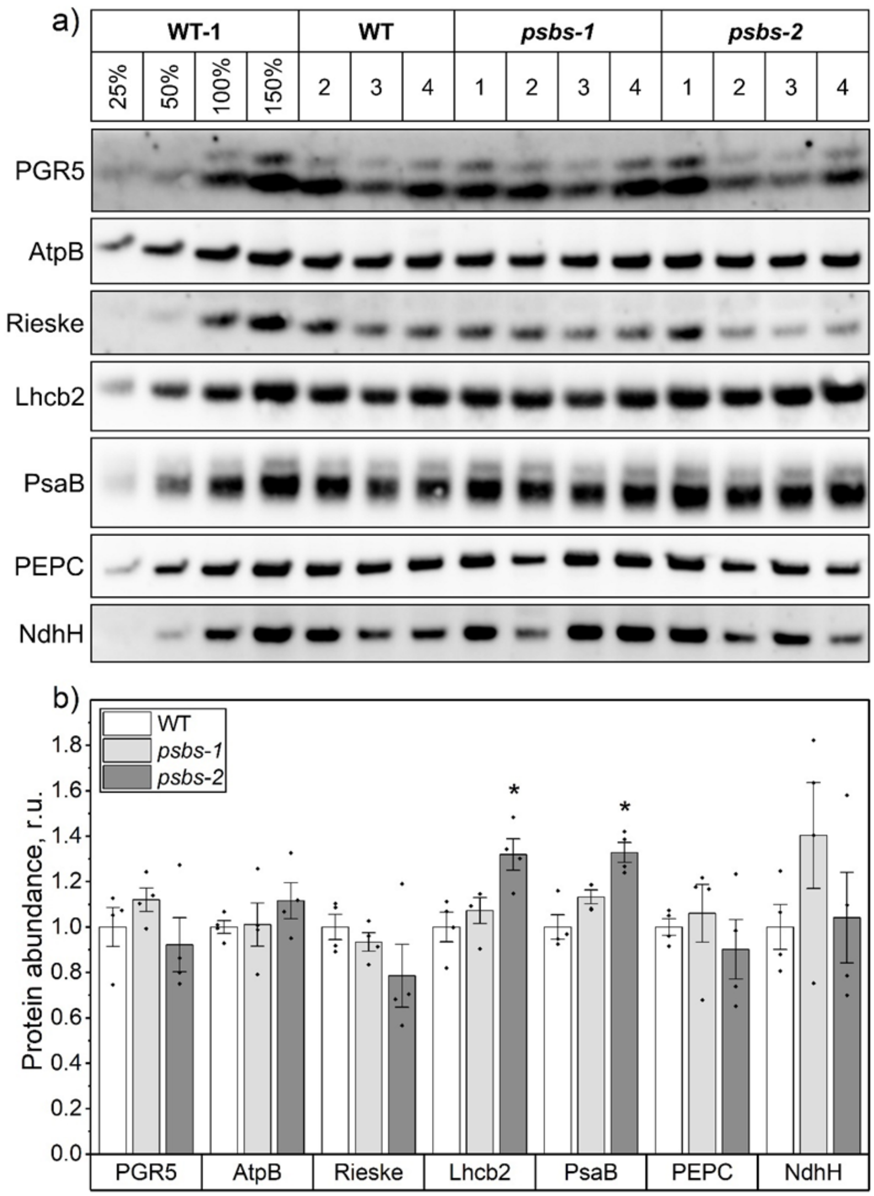


**Fig. S3.** Abundance of photosynthetic proteins in wild type (WT) *S. viridis* and plants lacking PsbS (*psbs-1* and *psbs-2*). (**a**) Immunodetection of PGR5, AtpB (ATP synthase), Rieske (Cytochrome $b_6f$), Lhcb2 (Light-harvesting complex II), PsaB (Photosystem I), PEPC, NdhH (NDH complex) in leaf protein samples isolated from plants grown under constant daylight and loaded on leaf area basis. (**b**) Relative quantification of protein abundance per leaf area. Mean ± SE with four biological replicates shown. Each protein has its own relative scale. Asterisks indicate statistically significant differences between PGR5-deficient and WT plants (1-Way ANOVA with Tukey's post hoc test at $P < 0.05$).

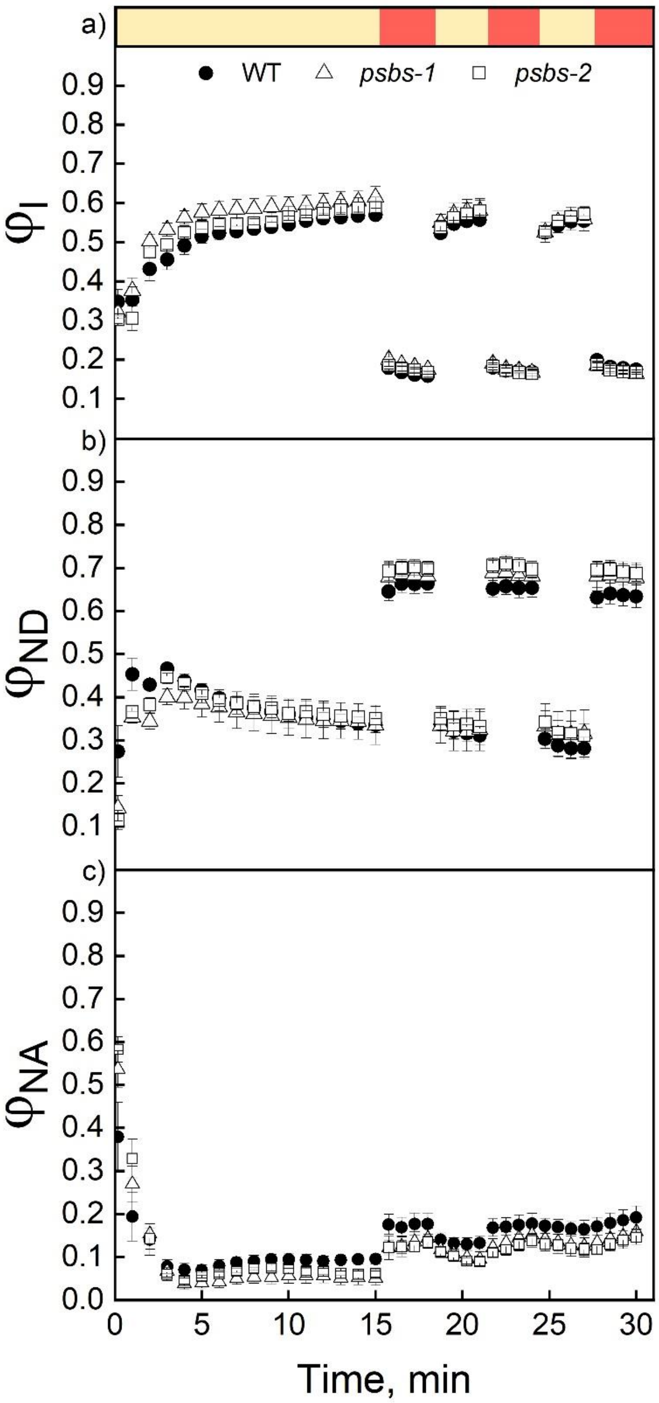


**Fig. S4**. Response of quantum yields within PSI to fluctuating light in wild-type (WT) *S. viridis* and plants lacking PsbS (*psbS-1* and *psbS-2*) grown in control conditions. (**a**) The photochemical yield of PSI ($\varphi_I$). (**b**) The non-photochemical yield of PSI due to donor side limitation ($\varphi_{ND}$). (**c**) The non-photochemical yield of PSI due to acceptor side limitation ($\varphi_{NA}$). Mean ± SE, $n$ = 5 biological replicates. No significant differences were found between genotypes (1-Way ANOVA with Tukey's post hoc test at $P < 0.05$).

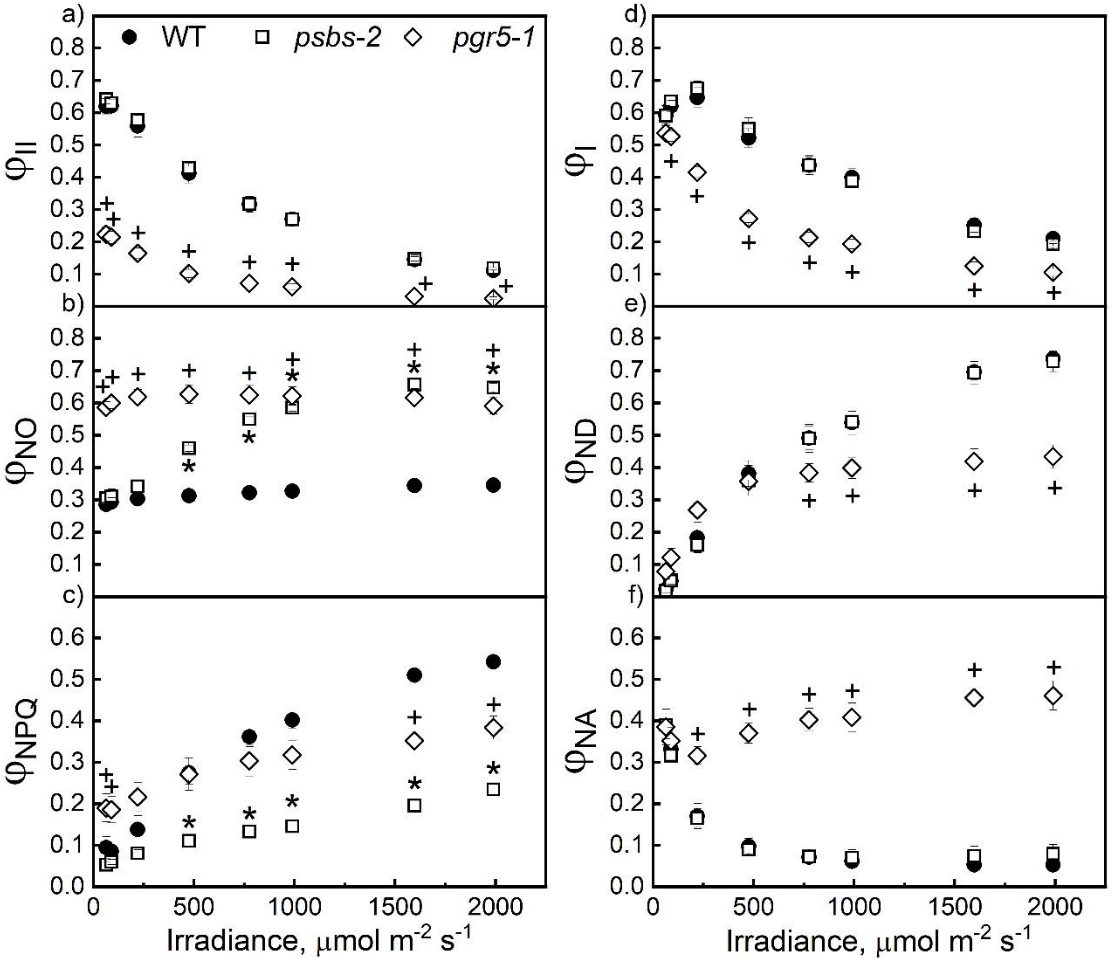


**Fig. S5.** Light response of quantum yields within PSII (**a** to **c**) and PSI (**d** to **f**) in wild-type (WT) *S. viridis* and plants lacking PsbS (*psbS-2*) or PGR5 (*pgr5*-1) grown at fluctuating daylight conditions (repeating cycles of 5 min at 250 µmol $m^{-2}$ $s^{-1}$ and 1 minute at 1000 µmol $m^{-2}$ $s^{-1}$). (**a**) The effective quantum yield of PSII ($\varphi_{II}$). (**b**) The yield of nonregulated non-photochemical reactions ($\varphi_{NO}$). (**c**) The yield of NPQ ($\varphi_{NPQ}$). (**d**) The photochemical yield of PSI ($\varphi_{I}$). (**e**) The non-photochemical yield of PSI due to donor side limitation ($\varphi_{ND}$). (**f**) The non-photochemical yield of PSI due to acceptor side limitation ($\varphi_{NA}$). Mean ± SE, *n* = 4 to 5 biological replicates. Asterisks indicate significant differences between WT and *psbs-2*. "+" symbols indicate significant differences between WT and *pgr5-1* (1-Way ANOVA with Tukey's post hoc test at $P < 0.05$).

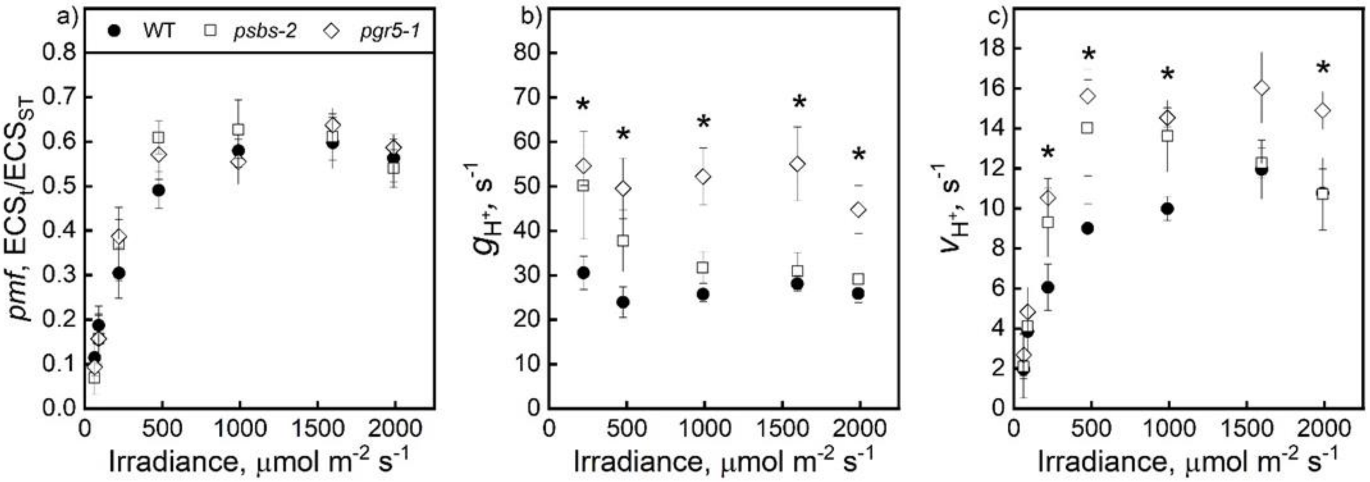


**Fig S6**. Light response of thylakoid membrane energisation parameters in wild-type (WT) *S. viridis* and plants lacking PsbS (*psbS-2*) or PGR5 (*pgr5-1*) grown at fluctuating daylight (repeating cycles of 5 min at 250 µmol $m^{-2}$ $s^{-1}$ and 1 minute at 1000 µmol $m^{-2}$ $s^{-1}$). **(a)** Proton motive force (*pmf*), **(b)** Proton conductivity of the thylakoid membrane ($g_{H}$+). **(c)** The light-driven proton flux across the thylakoid membrane ($v_{H}$+). Mean ± SE, *n* = 5 biological replicates. Asterisks indicate significant differences between WT and *pgr5-1*; no significant difference was found between *psbs-2* and WT (1-Way ANOVA with Tukey's post hoc test at $P < 0.05$).

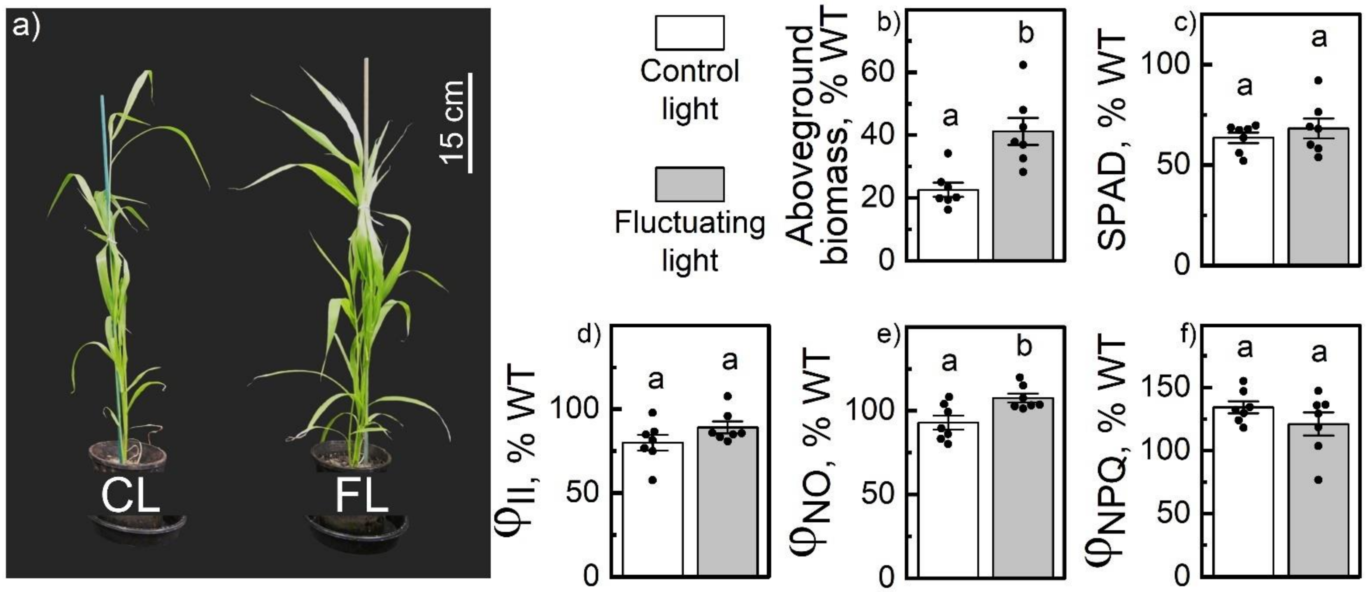


**Fig. S7**. Growth and photosynthesis of *S. viridis* plants lacking NDH complex (*ndhO-2*) grown either under control conditions (constant day irradiance of 380 $\mu$mol $m^{-2}$ $s^{-1}$; CL) or fluctuating daylight (repeating cycles of 5 min at 250 µmol $m^{-2}$ $s^{-1}$ and 1 minute at 1000 µmol $m^{-2}$ $s^{-1}$; FL). (**a**) Phenotype of plants 5 weeks after germination. (**b**) Aboveground biomass of plants at harvest. (**c**) Relative leaf chlorophyll content (SPAD). (**d**-**f**) Leaf photosynthesis parameters analyzed with MultispeQ at ambient light (at low light phase for fluctuating daylight): the effective quantum yield of PSII ($\varphi_{II}$), the yield of nonregulated non-photochemical reactions ($\varphi_{NO}$), and the yield of NPQ ($\varphi_{NPQ}$). (b-f) values are relative to WT values in corresponding condition, as presented in Fig. 1d and Fig. 4b for (b), and Fig. 1e and Fig. 4d for (c). For WT under constant light $\varphi_{II} = 0.45 \pm 0.01$, $\varphi_{NO} = 0.24 \pm 0.01$ and $\varphi_{NPQ} = 0.31 \pm 0.01$. For WT under fluctuating light $\varphi_{II} = 0.57 \pm 0.02$, $\varphi_{NO} = 0.21 \pm 0.01$ $\varphi_{NPQ} = 0.22 \pm 0.02$. Mean ± SE, $n = 7$ biological replicates. Letters indicate significant differences between groups (One-way ANOVA with Tukey's post hoc test at $P < 0.05$).
